# A unifying model to predict multiple object orienting behaviors in tethered flies

**DOI:** 10.1101/379651

**Authors:** Andreas Poehlmann, Sayan J. Soselisa, Lisa M. Fenk, Andrew D. Straw

## Abstract

Individual visual processing circuits for *Drosophila* locomotor control have been studied in detail, but contributions of specific pathways to multiple behaviors remain unclear. To address how both flexible and stereotyped visual object response behaviors potentially share neural circuit components, we investigated models of asymmetric motion responses. Such models have predicted that object fixation without explicit neural encoding of position is possible. Here we investigated what neural circuits and behaviors are consistent with such models. In behavioral experiments on tethered flying flies, we found close correspondence between T4/T5-neuron dependent turning responses to objects and model output for high frequency perturbations. Furthermore, we found that the model predicts key results from several published accounts of stereotyped object tracking. The concurrence of experiment and theory suggests a neural substrate and algorithmic basis for stereotyped object tracking and informs future studies of flexible visual behaviors and their neural bases.

## Introduction

Vision is a key element in the survival and behavior of many animals. For sensing objects and to estimate self-motion, many species rely heavily on visual input. Some behaviors like chasing and stabilization need to operate with minimal delay while other behaviors may be more flexible. With its comparatively small brain and precise neurogenetic tools, *Drosophila melanogaster* is a model well suited for studying how fast behavioral responses to objects are implemented on an algorithmic and neuronal level.

Two important visual behaviors in flies are the optomotor response to panoramic motion and turning in response to objects. The “optomotor response” is a behavior which serves to minimize wide-field visual motion with compensatory movements (Hassenstein and Reichardt, 1956) and is thought to stabilize gaze and locomotion. Building on work from other fly species, the past decade has used Drosophila to identify specific cell types and biophysical mechanisms associated with the motion detection operations thought to underly such optomotor behaviors. Direction selective responses to visual motion are thought to first arise in so-called T4 and T5 cells (Strother et al., 2017; Maisak et al., 2013). Optomotor responses in walking flies are strongly impaired when T4 and T5 neurons (local motion detectors) are genetically silenced (Bahl et al., 2013). These cells provide the main visual input to horizontal system (HS) cells (Schnell et al., 2012), wide-field motion sensitive neurons that were long thought to be involved in optomotor responses. Indeed, activation of HS cells leads to turns of the head and wings (Haikala et al., 2013), whereas inactivation of HS cells reduces optomotor responses of the head (Kim et al., 2017). On the basis of behavioral responses of insects to visual motion, Hassenstein and Reichardt proposed 60 years ago an influential correlator model for direction selective visual motion detection based on the enhancement of motion in the preferred direction (Hassenstein and Reichardt, 1956; Reichardt, 1987). Based on visual responses in rabbits, Barlow and Levick proposed an alternative model for motion detection that depends on the suppression of null-direction motion (Barlow and Levick, 1965). Recent and ongoing studies suggest hybrid models with preferred-direction enhancement and null-direction suppression are most consistent with accumulating results based on physiological recordings from T4 and T5 cells (Haag et al., 2016,2017; Leong et al., 2016; Strother et al., 2017; Gruntman et al., 2018). While hybrid models best describe the details of the cellular-level mechanisms involved in motion detection, simulations of behavior and neurons postsynaptic to T4 and T5, such as HS cells, are well fit by correlator type models over a wide range of conditions (Lindemann, 2005; Schnell et al., 2010; Fenk et al., 2014).

In contrast to our knowledge about the circuit mechanisms for wide-f¡eld optomotor response, another class of important visual behaviors, namely the responses to visual objects, is less well understood at a circuit and computational level. Object tracking behavior is a highly stereotyped active reorientation of the fly to keep vertical objects near the visual midline in the direction of travel. Object fixation behavior in walking flies does not require output from T4 and T5 neurons (Bahl et al., 2013). In tethered flight, however, object tracking in a figure ground discrimination task depends on intact T4/T5 neurons (Fenk et al., 2014). Since optomotor behavior relies on visual motion information, and object tracking behavior requires positional information in the visual stimulus, it is worth considering if the neural circuits mediating the two behaviors are distinct. Classical models, based on a linearized mathematical description of single degree of freedom tracking behavior dynamics, separated responses into two terms, one purely position-dependent and one motion-dependent (Poggio and Reichardt, 1973; Reichardt and Poggio, 1975). Some researchers interpreted these results in a way that position information and motion information may be computed by distinct neuronal circuits, but the original paper (see Poggio and Reichardt 1973) highlighted that a mathematical decomposition to facilitate analysis does not require that the neural circuits implement such a decomposition. More recently, the argument was raised that motion-dependent systems cannot account for object tracking because they are susceptible to low-frequency drift away from the tracked object (Fox et al., 2013). This interpretation is problematic, since it has been shown that asymmetric motion detector models based on properties of a single type of visual neuron are able to mediate the responses associated to the position-dependent term as well as the responses associated to the motion-dependent term (Poggio and Reichardt, 1973, 1981; Fenk et al., 2014). More recently, it was shown that the dynamics of saccades in magnetically tethered Drosophila are different for objects and backgrounds (Mongeau and Frye, 2017), suggesting again that these behaviors are mediated by distinct circuits.

Despite the robust and stereotyped nature of object tracking when tested in some conditions, neural and behavioral responses in Drosophila to objects are more variabile under other conditions. Depending on the size or shape of the object (Maimon et al., 2008), stimulus history (Neuser et al., 2008; Shiozaki and Kazama, 2017; Kuntz et al., 2017; Heisenberg and Wolf, 1984b), internal state (Gorostiza et al., 2016), the behavioral response to objects can also be more flexible (Heisenberg and Wolf, 1984b; Goetz, 1964). Furthermore, neural responses to objects have been described in central neuropils such as the optic glomeruli (Kim et al., 2015; Keleş and Frye, 2017) and central complex (Omoto et al., 2017; Shiozaki and Kazama, 2017; Sun et al., 2017; Seelig and Jayaraman, 2013, 2015) associated with higher-order behavior.

Therefore, here we compare predictions of visual-motor responses in a wide range of stimulus paradigms to those from a model based on asymmetric motion detection. To establish the validity of these predictions, we performed experiments to quantify behavioral responses of both intact flies and flies in which synaptic output from T4 and T5 cells was blocked with tetanus toxin and found a close correspondence between the T4/T5 dependent component of object responses and the model. We found that the model predicts, in open- and closed-loop simulations, many behaviors involving object tracking. Because this model is based on biologically plausible computations and is capable of predicting multiple behaviors, we suggest its utility in predicting some object tracking behaviors. We then discuss which other visual object behaviors are likely to depend on alternate mechanisms and circuitry. Thus, our work serves to illustrate what features of fly behavior can be predicted on the basis of current, biologically plausible models. Conversely, it shows what aspects of behavior remain unaccounted for. We suggest that objects responses with slower dynamics are not predicted by asmmetric motion detection models and may be indepdendent of T4/T5 circuitry.

## Results

### Perturbed object tracking of flies with intact and with blocked T4/T5 cells

We investigated the dynamics of T4/T5 dependent and independent neuronal circuits underlying object tracking behavior to test if these pathways contribute to behavior at different timescales. If one of these circuits mediates short timescale tracking behavior, blocking its input neurons will impair the fly’s ability to compensate fast perturbations in both the angular position and the angular velocity domain.

We ran tethered closed loop object tracking experiments on flies in which synaptic transmission was blocked in T4 and T5 neurons by expressing tetanus toxin (“T4/T5 block flies”, *;UAS-TNTe,tsh-GAL80;GMRSS00324-splitGAL4;*), and on flies that express an inactive variant of the toxin in the same neurons (“T4/T5 control flies”, *;UAS-TNTin,tsh-GAL80;GMRSS00324-splitGAL4;*). By using a highly specific split GAL4 driver line (Schilling and Borst, 2015), we minimized the possibility of motor defects resulting from off-target toxin expression. We used a tethered flight setup as depicted in Supplemental Figure 1A to show a visual stimulus that consisted of a 20deg wide black bar moving on a white background. The fly’s wing movement was analyzed using a video camera and realtime image processing software running at 120 Hz. The difference in leading edge wingstroke angles (delta wing beat amplitude Δ_WBA_) — approximately proportional to yaw torque (Tammero, 2004) — was measured in realtime and coupled back to give the fly control over its simulated orientation towards the bar (Supplemental Figure 1B). Closed loop single degree-of-freedom turning dynamics where approximated with a 2nd order differential equation as in Hesselberg and Lehmann (2007). Experiments consisted of closed loop fixation interrupted by 10s long trials during which a constant angular torque bias was added to the closed loop torque. Torques were randomly selected from a predefined set. To make comparison between perturbation torque and measured Δ_wba_ easier, torque is converted to its equivalent Δ_WBA_ perturbation offset throughout the following section.

Figure 1A shows averaged Δ_WBA_ for torque perturbation bias experiments in closed loop fixation. To continue fixating the bar, flies must compensate perturbation by adjusting their Δ_WBA_ to the given trial condition. Control flies were able to compensate the perturbation almost completely up to bias amplitudes slightly above 15 deg. T4/T5 blocked flies, on the other hand, show substantially decreased performance at lower perturbation strengths. Furthermore, they take several seconds to reach an average Δ_WBA_ plateau that is far below the required amplitude to compensate the perturbation and therefore fail to keep the stripe stationary. Investigating the plateau Δ_WBA_ reached by flies across varying perturbation offsets (Figure 1B) reveals that for very small offsets both blocked and control flies are able to adjust to the trial conditions (plateau amplitude on diagonal), but for absolute offsets larger than about 15 deg, blocked flies fail to adjust. The plateau Δ_WBA_ returns to almost zero for higher perturbation amplitudes. Δ_WBA_ distributions during non-perturbed bar tracking show that the maximum observed Δ_WBA_ range inT4/T5 blocked flies is larger compared to control flies (SupplementalFigure 1C), confirming that failure to compensate the perturbation is not due to a genetically introduced motor defect limiting maximum wing movement, but likely due to visual response reduction to the bar.

**Figure 1.**
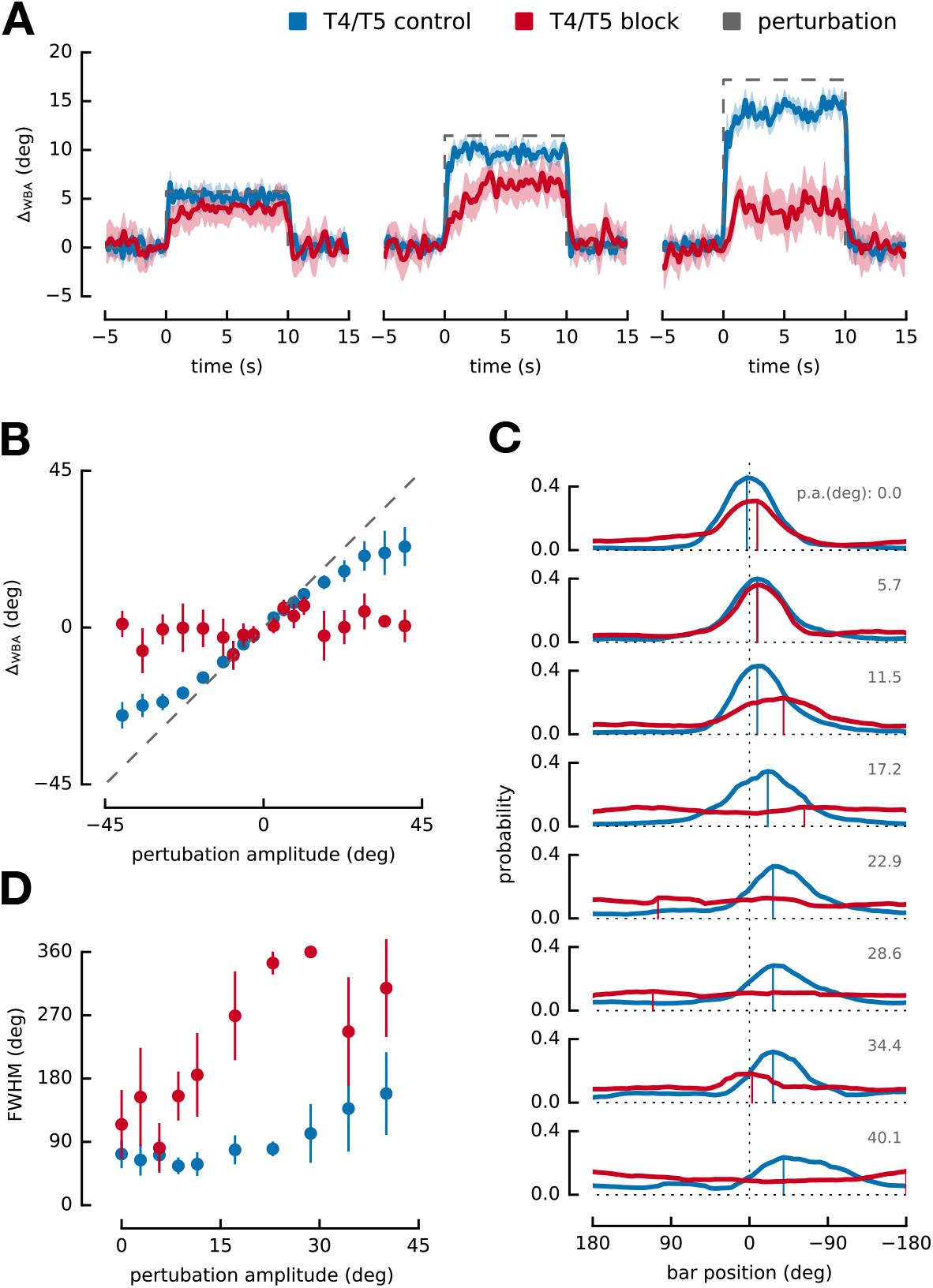
Delta wing beat amplitude offset experiment. Control flies blue, N=11, total number of trials m=55. Blocked flies red, N=9, m=45, (**A**) Averaged delta wing beat amplitude traces overtime for equidistantly spaced, increasing perturbation torque values. Shaded area indicates standard error of mean across flies. The dashed black line shows the Δ_WBA_ equivalent of perturbation torque over time for each trail. Control flies manage to almost fully compensate perturbation offsets up to roughly 15deg but show decreasing ability to do so at higher perturbations. Blocked flies already start to fail at perturbations offsets of about 10deg and take longer to plateau at a lower value. For increasing perturbation offsets blocked flies responses tend towards zero again. (**B**) Average plateau delta wing beat amplitude during last 5 s of perturbation. Dashed line indicates perturbation offset. Errorbars show standard deviation across individual flies. Control flies can compensate up to higher perturbation offsets. Blocked flies’ amplitudes tend towards zero for higher offsets. (**C**) Histograms of bar position for different perturbation offsets during last 5 s of trial. Negative offset amplitude trials have been mirrored and grouped together with positive amplitude trials. Top to bottom shows increasing perturbation amplitude offsets. Mode of distribution is marked with vertical line. At small offsets, distribution modes are shifted in same directions for control and blocked flies. (**D**) FWHM of bar position distributions during offset perturbation. Dataset mirrored and combined like in (C). Errorbars indicate standard deviation across flies. The FWHM increases dramatically for blocked flies with increasing perturbation amplitude beyond 15deg. Supplemental Figure 1. related to this.

Figure 1C shows distributions of angular position during the last 5 s of the trial duration, at which point transient changes in the position distribution are negligible. While the width and mode of the distributions are very similar between control and blocked flies during low amplitude trials, the distributions for blocked flies become relatively uniform for perturbation offsets larger than 11.5 deg. As a distribution-independent measure of fixation quality, we plot the full width at half maximum (FWHM) of angular position distributions during the last 5 s of the trial in Figure 1D. While blocked flies are able to keep the stripe comparably stationary at low perturbation amplitudes, they rapidly fail to do so with increasing perturbation amplitude. This would then explain why tracking responses in blocked flies are completely disrupted at lower perturbation offsets compared to control flies, and why measurements of positional responses comparing blocked and control flies without external perturbation should not differ in shape, but in amplitude (as seen in Bahl et al. 2013). −>

### Object tracking dynamics of flies with intact and with blocked T4/T5 cells

Because of the observation that the T4/T5 block results in an inability to compensate fast perturbations, we investigated the dynamics of object tracking. We used a similar method to the one previously described in Roth et al. (2012) to measure the open-loop system dynamics of object tracking under perturbed closed-loop conditions. It directly measures the transfer function describing the fly’s ability to track moving black vertical bars, assuming that the tethered flight tracking behavior can be well approximated by a linear time-invariant system. This assumption was shown to be valid within margins of error by Roth et al. 2012. We record closed-loop fly behavior while a time-varying perturbation is applied to the position of a black bar, which allows the gain (amplitude) and phase delay of the behavioral response to be estimated as a function of the perturbation frequency of the bar angle. The frequency dependence of this transfer function gives direct and intuitive insight into how well a fly is able to follow the trajectory of a moving bar and how much it lags behind the bar’s movement. As mentioned, the fly’s Δ_WBA_ is closed-loop coupled to the stimulus, so that the fly is in control over its orientation in the world coordinate system (see Supplemental Figure 2A). A black vertical bar of width 15 deg is centered at angular position 0 deg in world coordinates, and its position is perturbed during the experiment. The perturbation is defined by a logarithmic chirp of increasing frequency and decreasing amplitude (see Materials and Methods). This limits the maximum apparent angular velocity during the stimulus presentation, and still covers a wide range of perturbation frequencies, while avoiding to push the fly’s tracking behavior responses into a highly non-linear regime.

Figure 2A shows the averaged fly orientation in the world coordinate system as well as the Δ_WBA_ for control and blocked flies during the chirp stimulus. Initially, flies of both genotypes are able to change their orientations to follow the perturbation, and require only small modulation in Δ_WBA_ since the bar is moving comparatively slowly. Control flies keep the bar at the visual midline over the whole time course of the trial. Their orientation follows the trajectory defined by the logarithmic chirp and the Δ_WBA_ is most strongly modulated at around 60 s. At roughly 80s the fly does not completely follow every oscillation of the bar anymore, but still keeps it near the front. Blocked flies, on the other hand, lose the ability to reliably track the bar at around 40 s, as indicated by the FWHM of the bar position distribution reaching 360 deg (see Supplemental Figure 2B). Additionally, due to the loss of fixation, the stimulus progression had to be reset more often compared to control flies (refer to Materials and Methods and Supplemental Figure 2C). In the Δ_WBA_ time series, the blocked flies show an almost random response, fluctuating around zero, with large error. This could indicate that they can’t visually detect the bar anymore at these perturbation frequencies.

**Figure 2.**
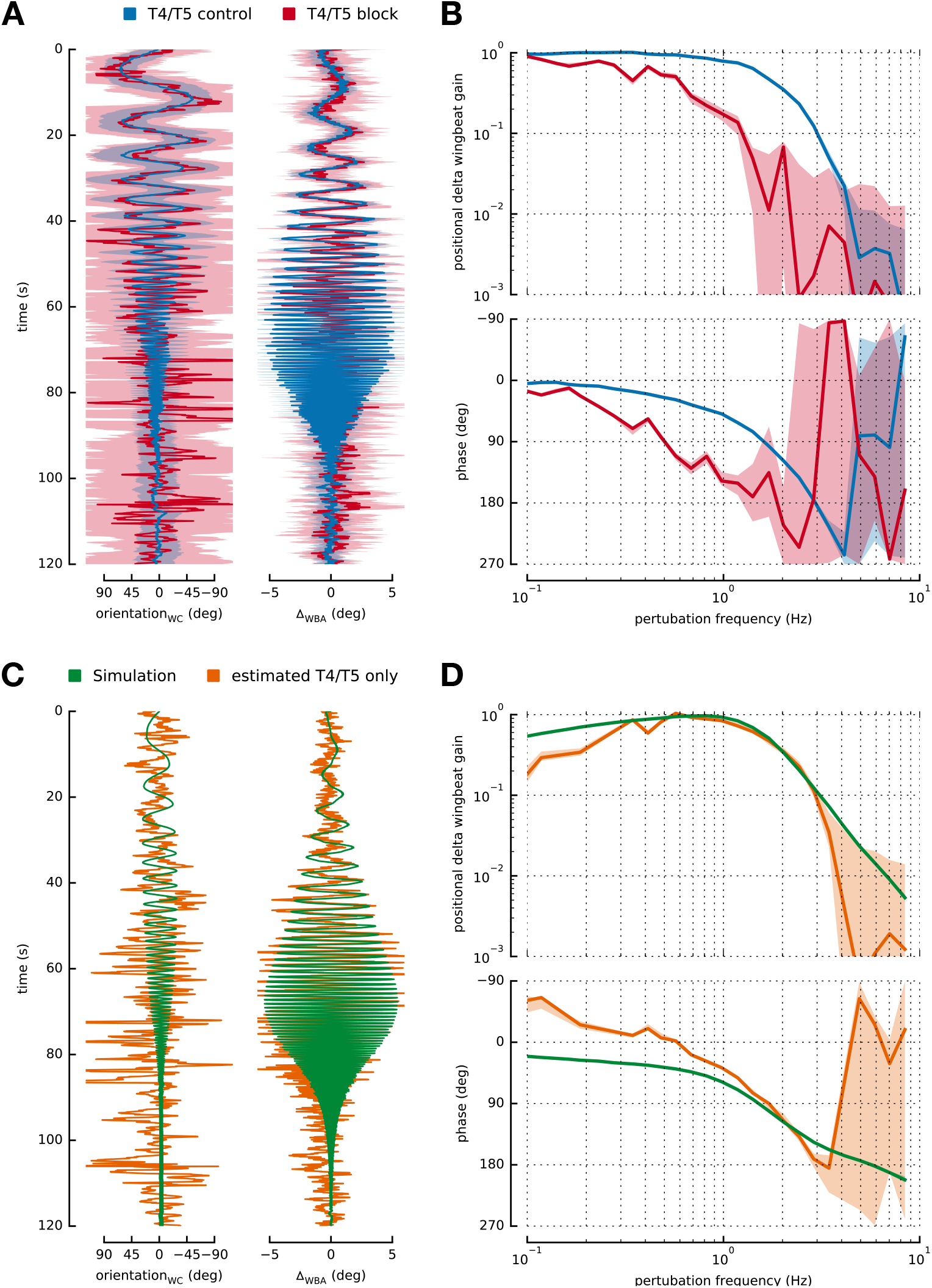
Logarithmic chirp experiment. Control flies blue, N=14, total number of trials m=66. Blocked flies red, N=14, m=64, (**A**) Averaged fly orientation in world coordinate system and median Δ_WBA_ over time. Shaded area shows FWHM for position plot and median absolute deviation for Δ_WBA_ plot. Position: Control flies follow the object position perturbation almost perfectly. Blocked flies do so for low perturbation frequencies but fail to orient correctly at higher frequencies. Δ_WBA_: While almost identical in performance at low perturbation frequencies, blocked flies fail to keep up with control flies starting at » 60s. (**B**) Bode plot of empirical transfer functions recovered from closed loop chirp experiments. Blocked flies tracking performance drops at lower frequencies compared to control flies. At 1 Hz blocked tracking performance is almost an order of magnitude worse than control performance. (**C**) As in (A), showing estimated T4/T5 only responses and numerical simulation results of an asymmetric motion system. Error estimates have been omitted for both. (**D**) As in (**B**). Approximated pure T4/T5 motion system responses should be interpreted as lower boundary on real transfer function. Above 0.6 Hz the approximated and numerical transfer functions are in remarkable agreement. Supplemental Figure 2. related to this.

Interpreting the chirp perturbation as the input to a linear system and the temporally integrated Δ_WBA_ as its output (using the closed-loop dynamic equation), we can calculate the empirical transfer function for both genotypes (see Materials and Methods). Figure 2B shows the recovered transfer functions for control and blocked flies, plotting both gain and phase. A gain of 1 indicates a perfect match to the input offset position amplitude and a phase of 0 indicates zero lag following of the bar. Control flies are able to follow the perturbations in the position domain up to frequencies of about 2 Hz, after which the gain drops dramatically. The flies are still fixating the bar even at higher frequency (average bar position is at the visual midline, see Supplemental Figure 2B), but do not follow the oscillations anymore. The measured empirical transfer function is in slight discrepancy with Roth et al. (2012) which measured a dip in object tracking performance at around 1 Hz. It is possible that this measured dip is an artefact of the shadow based Δ_WBA_ tracking method, because when observing the flies at perturbation frequencies of around 1 Hz we measure an almost threefold increase in leg extensions (Supplemental Figure 2D), consistent with T4/T5 mediated landing responses (Schilling and Borst, 2015). This leg extension behavior might interfere with shadow based measuring techniques, but can be dealt with in image based measuring techniques, like the one used here (see Materials and Methods and Supplemental Figure 1B). T4/T5 block flies are able to follow perturbations at low frequencies, but show a continuous drop in tracking performance starting at 0.6 Hz. The almost linear drop in phase starting at 0.2 Hz indicates a slow response time for the T4/T5 blocked tracking response, in line with the bad tracking performance at higher frequencies. At perturbation frequencies of 1 Hz, the blocked flies’ gain is almost an order of magnitude worse than in control flies. The higher the perturbation frequency, the less the blocked flies are able to keep the bar stationary. These results are consistent with the idea that T4/T5 independent object tracking is sufficient to fixate very slow moving objects and that faster tracking requires intact T4/T5 neurons. Another, not mutually exclusive, explanation for the remaining slow object responses is an incomplete block of T4/T5 cells with tetanus toxin. In this case, the contribution of T4/T5 dependent responses would be, at low frequencies, an underestimate and T4/T5 independent responses would be an overestimate. Given the complete loss of fixation capability at high frequencies upon T4/T5 block, we exclude the possibility of an T4/T5 independent system being required for fast tracking. Taken together, these experimental results suggest that T4/T5 independent object tracking system is only able to mediate black bar tracking up to perturbation frequencies of « 0.6 Hz, above which tracking behavior depends on T4/T5 dependent circuitry.

### Simulation of pure motion-vision based object tracking dynamics

In previous work, we showed that models based on asymmetric motion responses are able to mediate object fixation (Fenk et al., 2014). These models were based on correlator type motion detection, although the key property is the asymmetric motion response rather than the specific nature of the motion detection operation. The asymmetry refers specifically to the widely reported feature of several cell types postsynaptic to T4/T5 that the membrane potential depolarizes more to preferred direction motion than it hyperpolarizes in response to an equal magnitude opposite direction motion. Fixation arises when such model neurons are coupled to turning behavior because turns toward objects moving away from the midline are larger in amplitude than towards objects moving towards the visual midline. Fixation in these models arises as a dynamic equilibrium of the tendency to turn left and turn right being balanced at the midline. In the presence of background motion or biased feedback, the equilibrium shifts away from the midline, and this phenomenon is seen in models and behavioral experiments (Fenk et al., 2014).

We would like to compare the experimental results above on the dynamics of T4/T5 dependent behaviors with the predictions of biologically-plausible, visuo-motor models of fly T4/T5 dependent circuitry. Therefore, we tested our previously published computational model of the *Drosophila* visuo-motor system (see Materials and Methods) with the same logarithmic chirp perturbed closed-loop object tracking conditions as our real fly experiments. Ideally, we would want to compare the results of the simulations to responses of flies in which every visual response independent of the T4/T5 cells is blocked and thus to experimentally measure behavioral responses mediated by T4/T5 dependent circuitry alone. Since this is not currently technically possible with genetic or other means, we can consider the possibility that object tracking behavior across flies can be described as the linear sum of the T4/T5 independent and T4/T5 dependent systems. (Later we will consider non-linear summation.) Using this approach, we recover an estimation of the response due solely to the T4/T5 dependent pathway as the difference of the averaged control and blocked fly responses. Figure 2C shows fly orientation and Δ_WBA_ for the estimated “pure T4/T5 system” (orange) as well as the simulated asymmetric motion model (green). While it is hard to make any meaningful comparison in the position domain, the Δ_WBA_ results are in good agreement for higher perturbation frequencies. The empirical pure T4/T5 object tracking transfer function, as well as the simulated asymmetric motion model transfer function, are shown in Figure 2D. The experimental and simulation data are in remarkable agreement at frequencies above 0.6 Hz, which is to be expected if the two following points are true. First, the approximation of linear summation between the T4/T5 dependent and independent systems seems to hold. And second, that the computational model predicts the T4/T5 dependent behavioral responses well.

The discrepancy at lower frequencies (below 0.6 Hz) is consistent T4/T5 independent pathways being sufficient to compensate for slow perturbations. If the two object tracking systems mediate redundant object responses at low frequencies, it directly follows that estimates of the pure T4/T5 motion response depending on a linear summation assumption will always underestimate the real contribution of the T4/T5 motion system, because perfect closed loop fixation can never exceed a gain of unity. The possibility of incomplete block of the T4/T5 system at low frequencies would similarly result in underestimation of T4/T5 dependent behavioral responses. Thus, our estimated pure T4/T5 response here represents the lower bound of object tracking performance expected when all visual pathways except those that are T4/T5 dependent are blocked.

Overall, the experimentally approximated and numerically simulated pure T4/T5 system transfer functions agree at frequencies above 0.6 Hz, at which the influence of a T4/T5 independent system should be negligible. Interestingly, the simulated asymmetric motion system predicts very robust fixation at high frequency perturbations, which the blocked flies completely fail at, but control flies excel at. It should be noted that, since our computational model is based on a Hassenstein-Reichardt motion detector model, it shows no response in the limit of zero velocity. The measured transfer functions for blocked flies and control flies, together with the simulated asymmetric motion system transfer functions, suggest that black bar tracking responses may be redundantly mediated by a T4/T5 independent and T4/T5 dependent system up to frequencies of about 0.6 Hz. For higher perturbation frequencies, our results suggest that object tracking behavior is primarily mediated by the T4/T5 dependent motion-vision system.

### Prediction of closed loop object tracking tasks using motion-vision only

Due to the unimportance of T4/T5 independent circuits for high frequency fixation discussed above and the ability of our computational model to simulate the behavior of the T4/T5 dependent responses, we decided to simulate previously published object tracking experiments under various closed loop tethered flight conditions to retroactively test the “predictive” power of this model. All following experiments were originally done on flies with intact T4/T5 neurons, and some were used to justify or hypothesize the need for additional, non-motion based object response circuitry in the fly visual system based on the observed behavior. To be able to intuitively understand the results shown in this section it is best to think about these experiments in terms of the analytical description formulated in our previous work (Fenk et al., 2014). Briefly summarizing, given a narrow vertical object stimulus, a motion-vision based visual system model with asymmetric response amplitudes towards progressive and regressive motion can be mathematically described by a linearized differential equation, which separates into a position dependent term and a motion dependent term. The position dependent term is mainly defined by the difference in receptive fields of the inputs to the elementary motion detectors. The motion response is mainly defined by the delay time constants used in the correlator model. For a vertical bar stimulus on a structured background, the linearized second order differential equation can be expressed as:

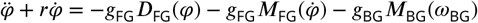

where *φ* describes the bar position relative towards the visual midline, *r* emulates the aerodynamic dampening, g_FG_ and g_BG_ are the closed loop figure and background gain, *D*_FG_ and *M*_FG_ are the position and motion response terms originating from the object and the asymmetric motion response, *M*_BG_ is the motion response term for the background, and ®_BG_ is the angular velocity of the background. Under normal gain coupling conditions *D*_FG_ yields a stable f¡xpoint at the visual midline. *M*_FG_ and *M*_BG_ (if moving together with the bar) render velocity 0deg s^−1^ a stable condition.

To facilitate direct comparison of our present simulations with the previously published experimental results, our figures have been made in a similar style as the original publication using only results from our simulations.

Monocular object tracking behavior has been previously measured in *Musca domestica* to investigate the influence of the binocular overlap region on the angular position dependence of object responses (Geiger et al., 1981). Figure 3A shows simulation results for black bar position histograms for simulated *Drosophila* under normal coupling and various eye occlusion conditions. From top to bottom, these conditions are “no occlusion”, “left eye covered with screen”, “right eye covered with screen” and “right eye covered with paint”. Since the background is unstructured, the only relevant term for tracking is the position dependent term *D*_FG_(*φ*). When no eye is occluded the bar is fixated at the visual midline, where *D*_FG_ has its zero crossing. When the screen was placed on one side, fixation was only possible when an additional torque offset was added to the torque produced by the simulated fly, and half of this offset torque was used to produce the covered eye results (compare with Geiger et al. 1981). These results are intuitively easy to understand when considering that the occlusion of one eye will basically set *D*_FG_ to zero on one half of the visual field, effectively removing its zero-crossing required for fixation. Adding a constant torque offset to the bar shifts the position response function vertically, creating a new zero-crossing along the edge of *D*_FG_, which, depending on its slope, is shifted away from the visual midline (see Fenk et al. 2014 for reference; in agreement with Virsik and Reichardt 1976). That covering an eye with paint creates a less steep slope of the fixation histogram is explained by the decreased slope at the visual midline of *D*_FG_ when only one receptive field is considered. Together with a shift of the onset of the position response during open loop rotation experiments (see Supplemental Figure 3), our model reproduces the tracking experiments from Geiger et al. (1981) using an asymmetric motion vision based model, without any motion-independent based position system.

**Figure 3.**
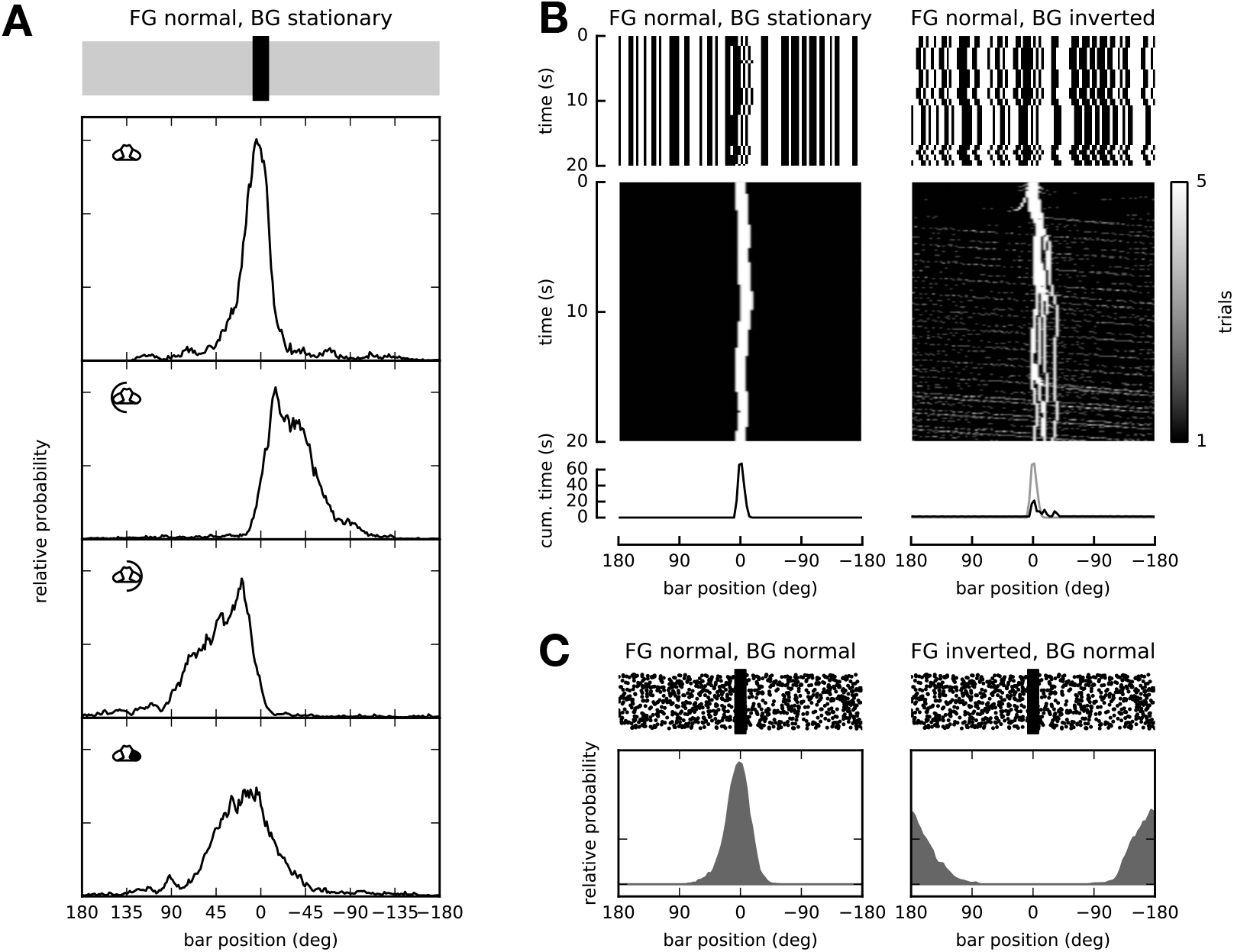
Closed loop simulation results. (**A**) Comparison of tracking behavior of flies under normal conditions (top), with a screen placed in front of the left eye with torque offset (2nd row), with a screen placed in front of the right eye with torque offset (3rd row) and the right eye covered with paint with torque offset (4th row). One eyed fixation distributions roughly double in width compared to normal fixation. Most likely fixation positions are shifted away from the occluded eye. See Geiger et al. (1981) Figure 6. (**B**) Inverted background gain simulation. Top row: space-time plots of stimuli. Middle row: space-time plot of tracking positions. Bottom row: histogram of cumulative time of bar at angular position. Left column: Bar coupled normally, background stationary. The bar is fixated in front of the fly across several trials, as can be seen by maximum of histogram. Right column: Bar coupled normally, background coupled inversely. While the stationary bar position at the visual midline is inherently unstable, it can stay there for some amount of time. The histogram shows a smaller but positionally consistent mode. See Fox et al. (2013) Figure 2. (**C**) Closed loop tracking of inversely coupled bar. Left: Background and bar are coupled normally and therefore move together. The bar is fixated at the visual midline. Right: Background coupled normally, bar coupled inversely. The stationary bar is still a stable condition but the stable position is now at the back of the fly. See Heisenberg and Wof (1984b) Figure 105. Supplemental Figure 3. related to this.

Inverted background gain coupling experiments have been previously used to hypothesize the active suppression of responses to moving wide field stimuli in the presence of a small field object (Fox et al., 2013). Figure 3B shows the result of an inverted background gain coupling simulation. The left column shows the simulation results for a stationary background. Across several trials, the bar is fixated at the visual midline. The right column then shows a rather artificial experimental scenario in which the torque generated by the fly is normally coupled to a small figure initially in front of the fly, but also inversely coupled to the background pattern. This means that movement of the bar produces opposite movement of the background pattern. It is argued that the reduced but non zero tracking behavior, as evident by the maximum of the bar position histograms, is proof that there must be a wide field background system and a figure system operating in parallel. Since g_FG_ is positive, the position in front and angular velocity zero would be a stable fixpoint. But the inverted background coupling together with the strong motion response towards the background renders the stationary bar inherently unstable. For very small deviations from the unstable position in front of the fly, the bar can stay roughly at the visual midline for a reasonable amount of time. If it deviates too much, the system locks into another stable behavioral mode, which is determined by the maximum torque produced by a spinning wide field pattern, and manifests as constant spinning of the background and the bar. To extend the time a bar can stay at the unstable point at the visual midline, we can reduce the closed loop gain to a weak coupling condition. Our simulation results reproduce this experiment with only a single pure wide field asymmetric motion system, however only under the condition of weak closed loop coupling compared to previous experiments (like in Figure 3A). Examination of maximum bar velocities (slopes of bar position traces) in Fox et al. (2013) suggest that these experiments were indeed run under weak coupling conditions (as indicated by the maximum observed slope of roughly 120 deg s^−1^). We predict that running similar experiments in which the initial bar position is at the side would reveal that the position at the visual midline is unstable.

Another artificial scenario which was used in learning experiments is inverted figure coupling with normally coupled background (Heisenberg and Wolf, 1984b). Figure 3C shows tracking behavior under these conditions. Again thinking about this in terms of the analytical model, this time the correctly coupled large background motion response, which outweighs the inversely coupled bar motion response, renders the zero velocity state stable. The inverted gain on the bar forces the positional term of the bar response to be unstable at the visual midline but now stable at the back of the fly. Our asymmetric motion model correctly predicts the initially measured anti-fixation behavior, but can of course not predict the learning behavior as shown in Heisenberg and Wolf (1984b).

In summary, our asymmetric motion based visual system model correctly predicts closed loop tracking behaviors with eye occlusion, under normal and inverted figure coupling as well as normal and inverted background coupling.

### Prediction of open loop object tracking tasks using motion-vision only

To further retroactively demonstrate the “predictive” power of our asymmetric motion model, we wanted to investigate different types of open loop behaviors that have been previously published.

Oscillating bar stimuli have been used to measure positional object responses because they reliably elicit visual tracking behavior (Maimon et al., 2008). Figure 4A shows the simulation of a fly responding to an open loop oscillating bar. The space-time plot (left) shows the time representation of the stimulus. A black bar of width 15 deg is oscillating at one of 5 different locations indicated by the black ticks on top. The Δ_WBA_ output of the simulated fly follows the oscillation of the bar. If the average position of the bar is to one side, the asymmetry in the motion response output leads to a shift of the average Δ_WBA_ to the same side. The right subplot shows the average Δ_WBA_ dependent on the average bar position. As it can be seen, this is in perfect agreement to predictions of the analytical model presented previously in Fenk et al. (2014) and basically shows the difference of the left and right receptive field of our visual model. The aversion behavior shown in Maimon et al. (2008) for black bars with smaller vertical extent can not be predicted by this one dimensional model.

**Figure 4.**
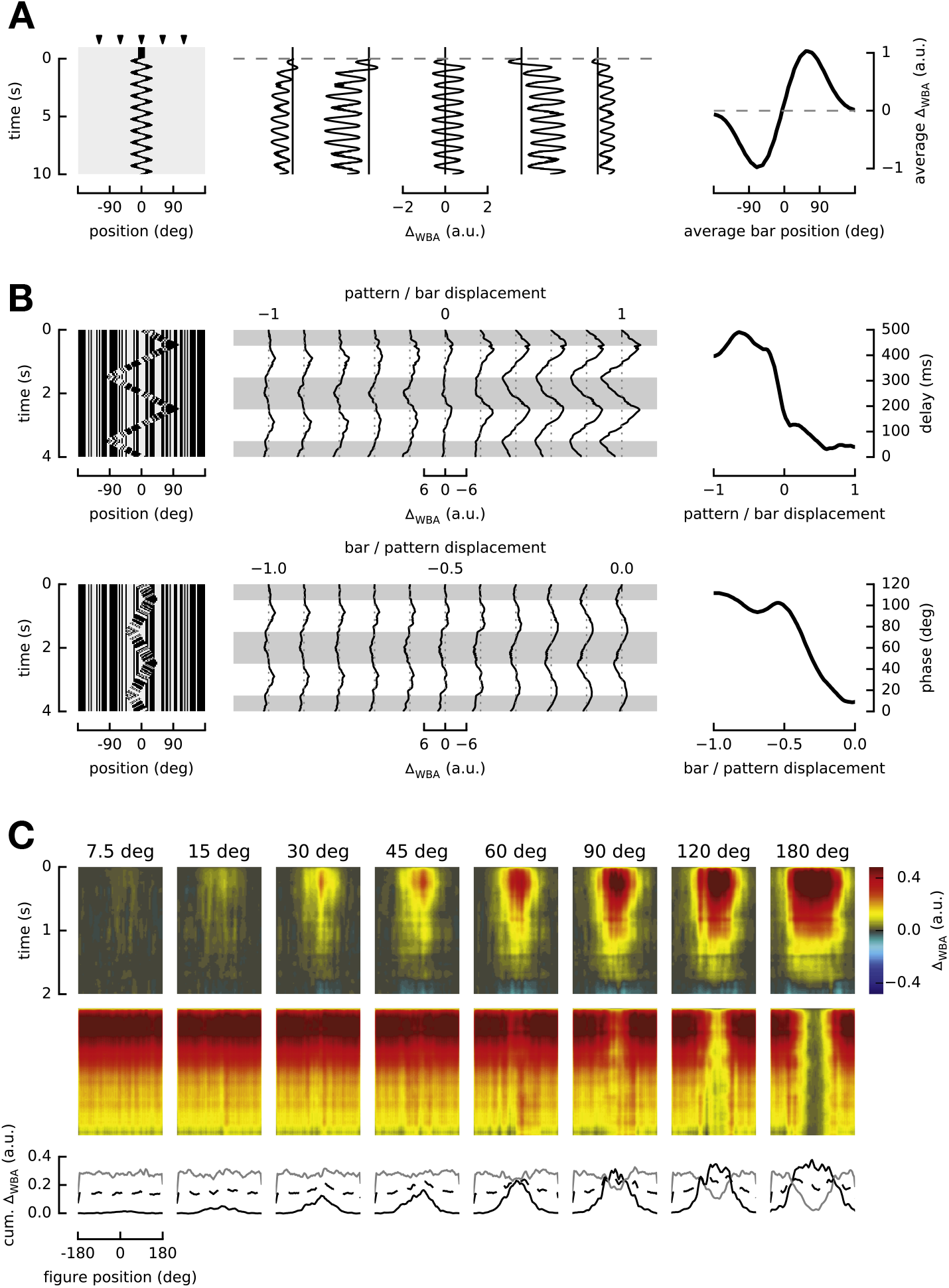
Open loop simulation results. (**A**) Δ_WBA_ responses to oscillating black bar of 15 deg width at different positions. Left: space-time representation of oscillating bar stimulus. Black ticks on top indicate the 5 different test conditions, at −120deg, −60deg, 0deg, 60deg and 120deg. Center: Δ_wba_ traces for the 5 tested stimuli. Δ_WBA_ shows position response like offset towards objects at either side. Right: Average Δ_WBA_ amplitude during trial for varying average bar positions. The resulting shape recovers the difference in receptive fields of the motion vision system. See Maimon et al. (2008) Figure 4. (B) Δ_WBA_ responses to theta motion stimuli. Left column: space-time representation of stimuli. Small bar is moving in a triangle wave pattern with periodicity of 2 s. Inner bar pattern is moving with same triangle wave function but 180deg phase shifted. Middle column: Δ_WBA_ traces for respective stimuli. Right column: delay and phase dependency on amplitude ratios between bar and pattern amplitudes. Top row: pattern/bar amplitude varying between −1 and 1. The delay in Δ_WBA_ responses ranges from roughly 500 ms down to sub 100 ms when varying the ratio. Bottom row: bar/pattern amplitude varying between −1 and 0. The phase shifts from ≈ 120deg down to sub 20deg when varying the ratio. See Theobald (2010) Figure 5/6. (C) Figure-Ground STAFs for varying figure widths. Top to bottom: Figure STAFs, Ground STAFs and averaged response of the first 100 ms of the STAFs. Left to right: Increasing figure width. The Figure STAF response becomes prominent enough at roughly 60deg to start to show a reduction in amplitude in the ground STAF. See Fox et al. (2013) Figure 6. Supplemental Figure 4. related to this.

Other more complex behaviors include responses to theta motion stimuli, which consist of moving objects which are themselves only defined by their internal pattern motion. Figure 4B shows simulation results of theta motion experiments like in Theobald (2010). The top row leaves the bar position amplitude fixed, while varying the bar/pattern amplitude ratio from −1 to 1. The bottom row leaves the pattern position amplitude fixed, while varying the pattern/bar amplitude ratio from −1 to 0 (refer to Theobald (2010) Figure 5/6 for space-time representations of each stimulus). The authors argued that the decrease in delay time across varying pattern/bar displacements from roughly 500 ms down to sub 100 ms, as well as the decrease in phase from roughly 120 deg down to sub 20 deg for varying bar/pattern displacement is a direct manifestation of the superposition of the outputs of separate subsystems with different dynamical properties. These subsystems would separately encode the 1st order motion of the object itself and the pattern motion inside the object. One of these systems would have short latency response and the other one a long latency response. We disagree with this argument because our asymmetric motion vision based model is able to reproduce this change in delay and phase without implementing multiple subsystems.

**Figure 5.**
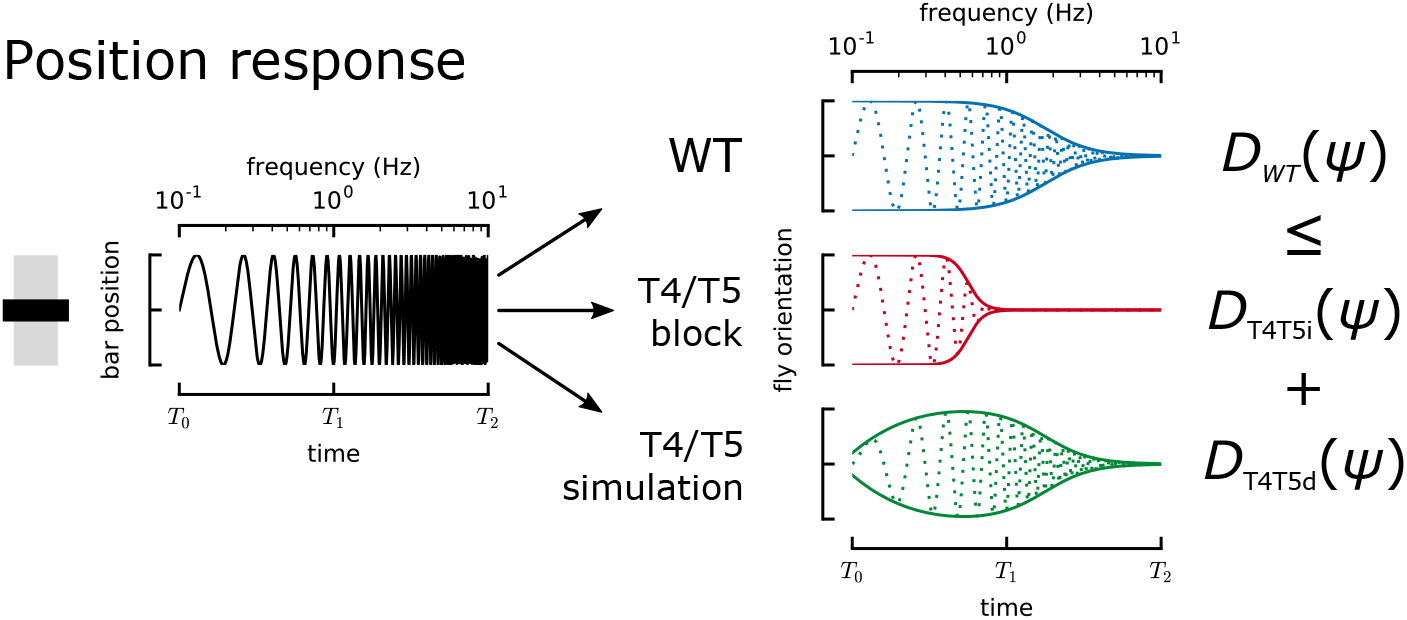
Summary: object tracking responses of a black vertical bar are mediated by T4/T5 independent (T4T5i) and T4/T5 dependent (T4T5d) circuits. Position responses are redundantly mediated by both systems at low perturbation frequencies. During short time scale perturbation, important during flight, only motion-computation based (T4/T5 dependent) systems are able to mediate object tracking. D is the position response component of fly turning behavior.

Finally, we wanted to investigate tracking behavior under a figure-ground discrimination experimental paradigm. Figure 4C shows results from spatio temporal action field (STAF) simulations as presented in Fox et al. (2013). STAF experiments simultaneously measure the impulse response towards two separate stepwise-moving patterns. One small-field figure pattern occludes the other wide-field background pattern. Impulse responses are then measured dependent on the position of the figure pattern during the measurement. The figure STAFs represent responses towards the figure pattern. Ground STAFs represent responses towards the background pattern. The figure and ground STAFs for increasing figure width show that starting at around 60 deg figure width we see a decrease in the amplitude of the ground STAF for positions in front of the fly. This becomes most evident when increasing the figure width even further. It has been argued that an active suppression of the ground response is responsible for the observed decrease in the ground STAF. This hinges on an observation that, when replacing the active figure with a gray pattern of the same width moving together with the background pattern, the suppression in the ground STAF does not happen. Our asymmetric motion based visual system model, with only a single motion vision pathway, produces similar STAFs showing “active” ground suppression for figures of widths above 90 deg (Figure 4C) and displays the lack of ground suppression for the gray occlusion experiments (Supplemental Figure 4). The model does not produce strong figure responses for very small figure widths, likely due to the wide receptive field width used in the model. Together, these STAF simulations show that a Hassenstein-Reichardt correlator based model with only an asymmetric motion response pathway shows figure responses and active background suppression without having any additional circuitry to implement such responses. We therefore disagree with the conclusion of Fox et al. (2013) that the STAF results argue that a position system is required to explain their key findings. While not all properties of the figure and ground STAFs can be recovered by simulating the experiments with these pure motion system models, the key aspects can be and thus our simulation illustrates that complex inputs to a simple system can generate complex outputs, without the need of active suppression mechanisms or parallel visual streams.

## Discussion

With our tethered flight behavioral experiments, we measured transfer functions for turning in response to visual object motion in flies with intact and genetically blocked T4 and T5 cells. The transfer functions show that motion-blind T4/T5-block flies are able to fixate an object that is perturbed at frequencies below 0.6 Hz, but fail to do so at higher perturbation frequencies. Additionally, we show that the difference in object tracking capability between control and blocked flies is well approximated using a Hassenstein-Reichardt-type elementary motion detector model with asymmetric motion response. This suggests that while object tracking can be mediated by a T4/T5 independent system for slow moving objects, it requires T4/T5 input if the object moves faster. We hypothesize that during free flight, where response times are critical, tracking behavior is predominantly mediated by a T4/T5 dependent motion-vision based circuitry (see Figure 5).

Additionally, we used the asymmetric motion visual system model to predict multiple different types of closed loop and open loop fly behavior, ranging from simple tracking behavior, across more complex anti-fixation behavior to theta motion responses and figure-ground discrimination. Because this relatively simple and biologically relevant model is able to predict this variety of experimental results without requiring adjustment of internal parameters for each specific condition, we suggest that this model is relevant for understanding the neural basis of these multiple behaviors. We know of no other model which is capable of predicting such a wide range of fly visuo-motor behaviors.

We suggest that future studies of object tracking behaviors need to test if these behaviors require T4/T5 based motion computation circuits, and if an impairment of the ability to track objects can be explained by a lack of asymmetric motion computation based object responses. We showed that many object behaviors (Geiger et al., 1981; Theobald, 2010; Fox et al., 2013; Virsik and Reichardt, 1976; Maimon et al., 2008; Heisenberg and Wolf, 1984a) can be explained by asymmetric models of T4/T5 dependent circuitry. Given the mathematical equivalence of the asymmetric motion dependent model with Poggio and Reichardt (1973), and its ability to compensate perturbations of the object position (see Figure 2 and Fenk et al. 2014), we predict object behavior during flight (like the chasing behavior measured in Land and Collett 1974), where dynamics are inherently faster, will depend on motion vision. This agrees with conclusions drawn in Reichardt and Poggio (1975) and fits with the observations made by Land and Collett (1974) that simulated chasing trajectories require velocity dependent damping terms. These velocity dependent responses help the fly initiate turns in the direction of the object motion even if this turning direction is opposite to a purely position dependent turning response, allowing the fly to do “predictive” object tracking (Land and Collett, 1974). Our transfer function measurements suggest that fast object tracking behavior will mainly be mediated by an asymmetric motion computation circuit, instead of slower T4/T5 independent object tracking circuitry (see Figure 2B). While our results highlight the capability of one pathway, they do not exclude the possibility of other pathways. When candidate circuit elements in the T4/T5 independent object tracking circuitry are genetically accessible, studies of chasing behavior in a silenced background will be able investigate directly the relative contributions.

The argument that pure visual motion based systems are ill-suited for object tracking because they fail to compensate low-frequency drift (Fox et al., 2013) cannot be accepted, especially when many of the stimuli presented to measure “position responses” contain motion-information at temporal frequencies above 0.6 Hz. When considering response behaviors, we suggest to use our parsimonious model to see if they can be predicted by an asymmetric motion system. Even seemingly complex position responses can be mediated by motion vision based models as long as they have a key property of asymmetry (a difference in response amplitude for progressive and regressive motion).

Despite the agreement of model predictions and some behavioral results presented here, the range of applicability of the model is limited. Several studies report behaviors which cannot be described in the present framework. For example, saccades towards objects have different dynamics than turns in response to background motion (Mongeau and Frye, 2017), but our modeling assumes a fixed relation between visual output and motor response. Neurons within central complex have been recently shown to maintain an internal representation of the fly’s heading during walking, both in the presence or absence of visual cues Seelig and Jayaraman (2015); Green et al. (2017); Turner-Evans et al. (2017). The same neuronal structure was also shown to be involved in visual memory tasks Neuser et al. (2008); Ofstad et al. (2011); Shiozaki and Kazama (2017). Recent work has furthermore shown that the anterior visual pathway (AVP) (Omoto et al., 2017) contributes to the recognition of novel objects (Sun et al., 2017; Shiozaki and Kazama, 2017), as well as suppressing contralateral stimuli. While the timescales required for such behaviors have not been investigated in detail, the timescales involved appear slower than those that require T4 and T5 cells. Further, these behavior experiments were performed in conditions beyond the range of validity of our asymmetric motion model by involving, for example, the role of prior experience or multiple motor systems with variable dynamics. It will be interesting to disentangle the relative contribution of these higher brain centers and purely motion-selective pathways to a fly’s orientation in space.

As a complement to future work investigating flexible object responses such as circuits in the central complex, models of object fixation mediated by asymmetric motion detection and consistent with results from the T4/T5 blocked flies will be valuable for clarifying the details of how particular neural structures are involved with specific behaviors.

## Acknowledgements

The authors thank Dan Bath, John Stowers and Abina Boesjes for discussion. This work was supported by European Research Council (ERC) starting grant 281884 and Wiener Wissenschafts-, Forschungs-und Technologiefonds (WWTF) grant CS2011-029 to A.D.S.

## Materials and Methods

### Flies

We used flies raised on yeast food at 25 °C. Female flies 4 days past eclosion were used for experiments. All experimental flies were the heterozygous F1 offspring of *GMRSS00324-splitGAL4* males (*w−;R59E08-AD;R42F06-DBD;) (Schilling and Borst, 2015*) and females carrying *UAS-TNTe* (active) or *UAS-TNTin* (inactive) and *teeshirt-GAL80* to block *GAL4* expression in the ventral nerve cord and peripheral neurons (*w-;UAS-TNTe,tsh-GAL80;+*;) or (*w-;UAS-TNTin,tsh-GAL80;+*;). Each fly was cold anesthetized for tethering and given a 30 min recovery period before being exposed to visual stimuli.

### Tethered flight setup

Experiments where done in a custom built tethered flight rig similar to the one used in Fenk et al. (2014) but with a cylindrical screen with 10cm diameter and 12cm height. The LED DLP projector used was an Optoma ML500 operating at 120 Hz. The closed loop latency of the system from delta wing beat amplitude change to photon was measured as 28 ms. The wing beat tracker runs at 120 Hz and is robust against partial occlusion due to leg extensions even if they are intersecting with the leading wing edge on the tracking image (Supplemental Figure 1B). As measured in Hesselberg and Lehmann (2007), the fly is given closed-loop control over its orientation *φ* via:

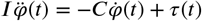

Here, the rotational inertia *I* is defined to be 0.52 × 10^−12^ N m s^2^. Aerodynamic rotational dampening *C* is set to 54 × 10^−12^ N m s. The estimated torque *τ* generated by the fly is calculated from delta wing beat amplitude Δ_WBA_ via:

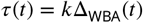

with *k* = 2.9 × 10^−10^ N m deg^−1^.

### Torque bias stimulus

The torque bias stimulus consisted of unbiased and biased closed loop object tracking trials, in which a 20 deg wide black bar could be tracked by the fly. The feedback loop was modulated during 10 s perturbation trials in which a constant angular torque offset was added, which is equivalent to adding a constant Δ_WBA_ offset to the measured Δ_WBA_. Since the feedback dynamics are dominated by aerodynamic dampening, the constant torque offset effectively manifests as an angular velocity bias on the bar. Each trial was triggered after the fly fixated the bar for at least 2 s. The minimum waiting time between two trials was 10 s. Torque ranges were initially determined so that we could observe T4/T5 blocked flies failing to track the bar. During experimental trials, preselected torques were randomized and tested in both directions.

### Chirp stimulus

The chirp stimulus consisted of a 15 deg wide black bar moving at a predefined trajectory in world coordinates, identical to Roth et al. (2012). The trajectory was given by a logarithmic chirp ranging from 0.05 Hz to 12.5 Hz over the time period of 120 s. The chirp amplitude *A* was modulated by A(ω) = A_0_ (0.918 + 0.264 s · ω)^−1^, dependent on the angular frequency *ω* and the initial amplitude A_0_ = 60 deg. The fly was in active control over its orientation in the world coordinate system, so that it could track the bar. To get reliable data for T4/T5 blocked flies, we had to interrupt the stimulus if flies were locked in a behavior that would spin the bar around. If the fly lost track of the object (indicated by the bar spinning around the fly more than 5 times per 5 s) we stopped the closed loop stimulus, reset the bar position to the visual midline of the fly for 5 s and returned closed loop control to the fly at the time when the first spin occurred. While this was rarely necessary for the control flies, it was required to get reliable data for the blocked flies (see Supplemental Figure 2C).

### Transfer function extraction

To recover the transfer functions from closed loop trials, we treat the tethered flight setup and the fly as a input-output system, where the perturbation signal over time is the input and the temporally integrated Δ_WBA_ is the output of the system. This includes the dynamics of the setup in the transfer function, but helps to improve the accuracy of the method for blocked flies, which fail to reliably track the bar across most of the trial. Since the setup dynamics only show low pass filter characteristics, and are dominated by the processing and display delay of 28 ms corresponding to « 35 Hz they can be ignored for the 12.5 Hz limited responses measured here. Δ_WBA_ is integrated via the closed-loop coupling dynamic equations described in the setup section. To recover gain and phase of the transfer function, the integrated output signal is divided into overlapping intervals in the time domain which are equidistantly spaced in the frequency domain. Each interval is then used to estimate amplitude and phase of the logarithmic chirp input signal using the output signal data. To estimate amplitude and phase we use a Bayesian estimation routine based on a Markov-Chain-Monte-Carlo method. Priors are chosen to be completely uninformative. Sampling is done until posteriors are reasonably well converged. Best estimates of actual gain and phase are chosen as the maximum likelihood values from resulting posterior distributions. Error estimates are 90 % confidence interval ranges calculated from posterior distributions.

### Numerical modelling

All software used for modeling will be released under an open source licence upon paper acceptance and made available at github.com. Numerical simulations use a Hassenstein-Reichardt-type elementary motion detector based visual system model, previously described in Lindemann (2005) with an asymmetric motion response (Fenk et al., 2014). The model was reduced to one dimension with inter-ommatidial angles of 5 deg and receptive fields matching horizontal system cell responses at 0deg elevation from Schnell et al. (2010). The *Calliphora* adequate LMC input filters were replaced with plain first order high-pass filters with a characteristic time of 80 ms. Time constants in first order low and high pass filters in both arms of the correlators were set to 140 ms to comply with results from *Drosophila* behavioral experiments (Leonhardt et al., 2016). Scaled model output is interpreted as Δ_WBA_ and limited via a first order low pass filter to a maximum change rate of 60 deg /s to emulate maximum observed change rate of Δ_WBA_ in control flies. Closed-loop simulations emulate the 28 ms tethered rig setup delay. Simulated experiments try to emulate high resolutions screens or LED cylinders dependent on experiment. Internal model parameters are identical across all simulations if not explicitly mentioned otherwise.

---

**Supplemental Figure 1.**
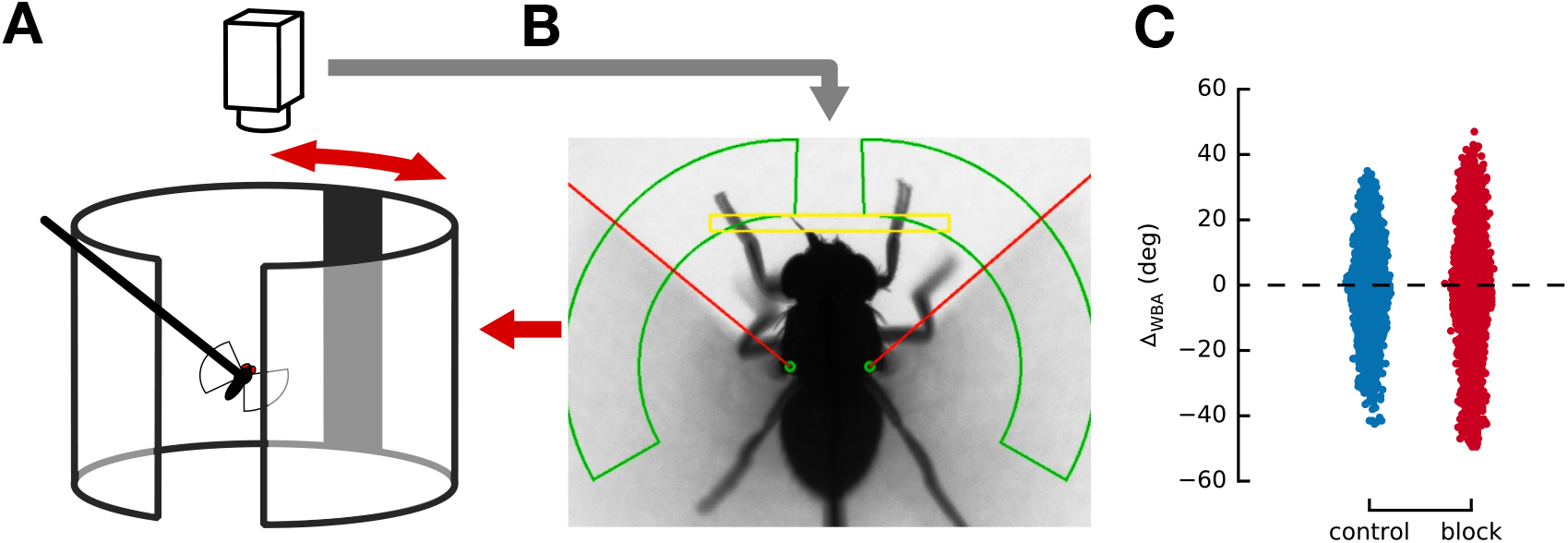
(**A**) Closed loop tethered flight setup. The fly is mounted on a thin tungsten tether in the center of a cylindrical projection screen. The visual stimulus is projected onto the screen with a DLP projector. A camera mounted above the fly is tracking the wing movement of the fly. Realtime analysis of delta wing beat amplitude allows for closed loop coupling with stimulus. (**B**) View of camera image used for realtime tracking of Δ_WBA_. Green framed areas indicate image portion in which the leading wing edge is tracked (detected edge marked by red line). Yellow framed area indicates image portion used to determine if the fly extends it’s legs. Leg extensions do not interfere with leading edge tracking of wing stroke in this image based method, but could interfere in shadow based methods. (**C**) Scatter plot of observed delta wing beat amplitudes during object fixation. The blocked flies show a broader distribution compared to the control flies.

**Supplemental Figure 2.**
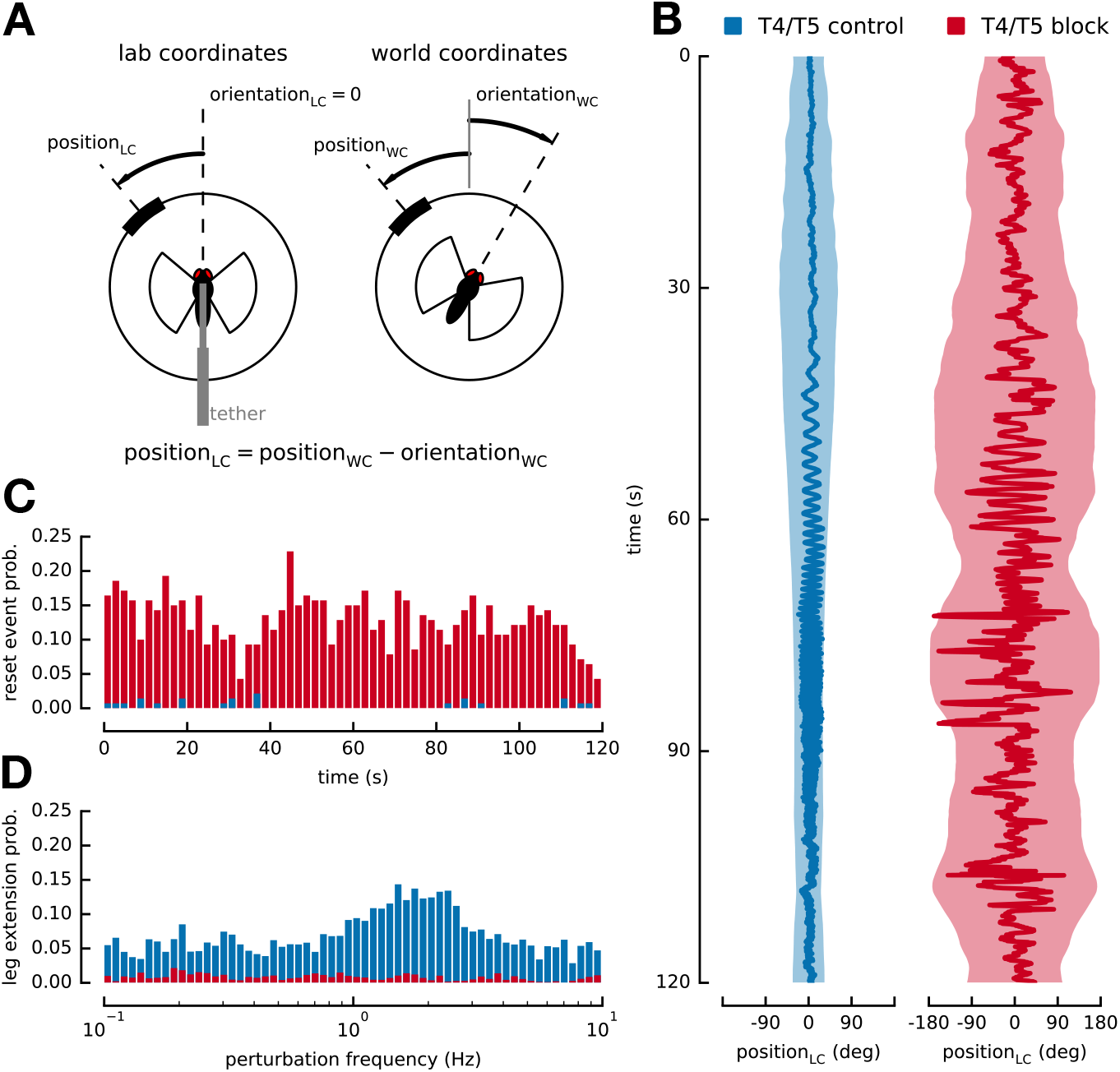
(**A**) Lab and world coordinates during chirp stimulus explained. (**B**) Filtered average bar position in lab coordinate system overtime. Shaded area marks FWHM of tracking distribution. Control flies fixate the bar almost perfectly. Blocked flies lose their ability to track reliably at around 45 s, indicated by a FWHM of almost 360 deg. (**C**) Reset event probability over time during chirp event. When the bar spins more that 3 times in 5 s, the stimulus is reset, to a stationary bar in front of the fly for 5 s, after which the stimulus resumes at a time prior to ‘time of first spin’ minus 5 s (see Materials and Methods). Stimulus interruptions are almost absent in control flies. Blocked flies average at a probability of about 10 %. (**D**) Leg extension event probability over time during chirp experiment. Blocked flies show almost no leg extension behavior. Control flies show an increased amount of leg extension between perturbation frequencies of 1 Hz and 3 Hz.

**Supplemental Figure 3.**
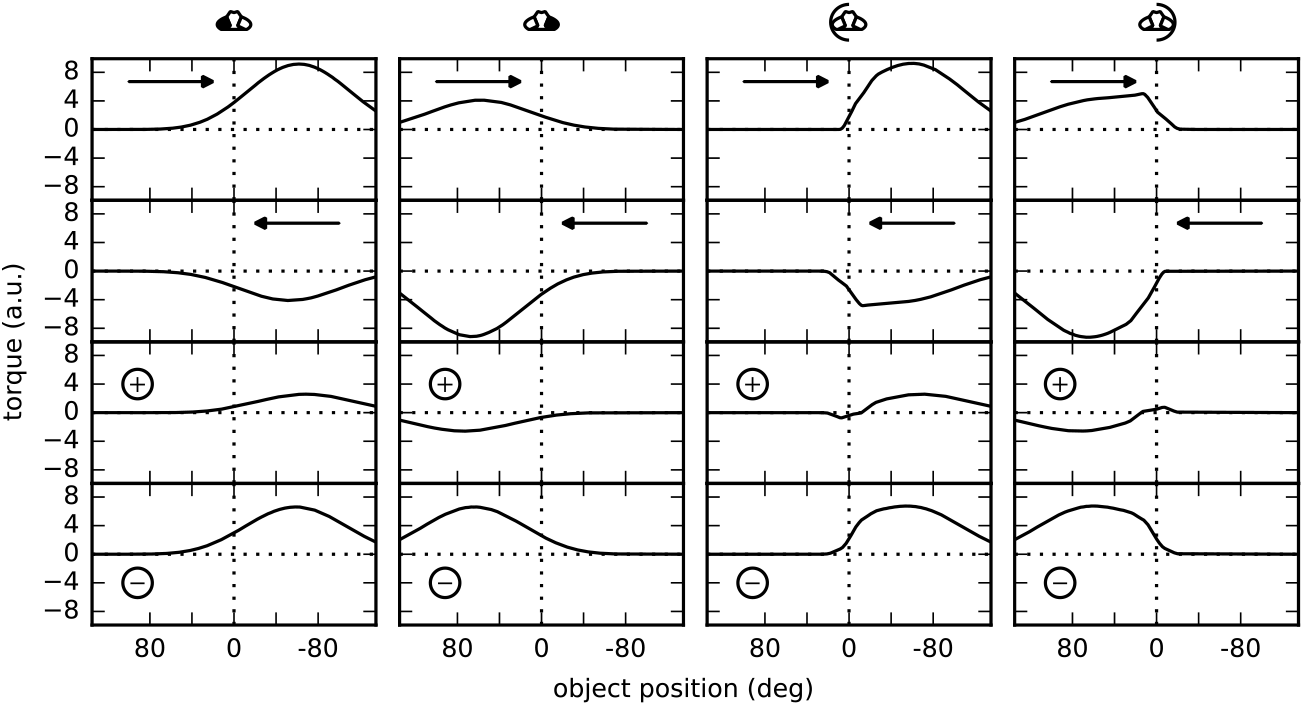
Simulated torque responses toward a moving black bar for flies where half of the visual field was occluded by a screen and for flies where one eye was covered with paint. Top two rows show the response to a bar moving from left to right and right to left respectively. Bottom rows show sum (position response) and difference (motion response) of the two top rows. Position responses and motion responses created by summation and subtraction qualitatively agree with experimental results. See Geiger et al. (1981) Figure 5.

**Supplemental Figure 4.**
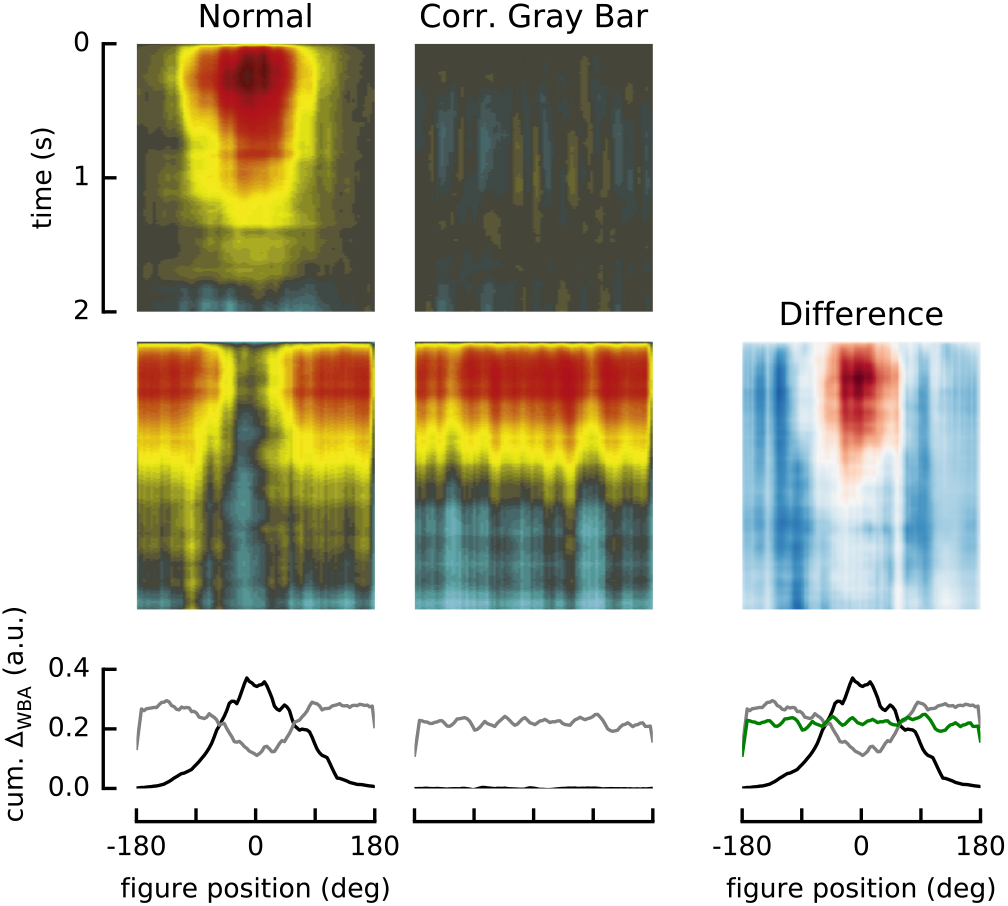
Control simulation for background suppression argument. Top to bottom: Figure STAFs, Ground STAFs and averaged response of the first 100 ms of the STAFs. The left column shows a normal experiment that shows background suppression with background pattern and uncorrelated figure. The center column shows an experiment in which the figure was replaced with a gray bar of identical width which moves correlated together with the background. The suppression in the ground STAF response vanishes vanishes when using a correlated gray bar as figure. The right column is plotted to facilitate easy comparison. See Fox et al. (2013) Figure 5.

